# Activity budget and gut microbiota stability and flexibility across reproductive states in wild capuchin monkeys in a seasonal biome

**DOI:** 10.1101/2021.08.09.455561

**Authors:** Shasta E. Webb, Joseph D. Orkin, Rachel E. Williamson, Amanda D. Melin

## Abstract

Energy demands associated with pregnancy and lactation are significant forces in mammalian evolution. To mitigate increased energy costs associated with reproduction, female mammals have evolved behavioural and physiological responses. Some species alter activity to conserve energy during pregnancy and lactation, while others experience changes in metabolism and fat deposition. Restructuring of gut microbiota with shifting reproductive states may also help females increase energy harvest from foods, especially during pregnancy. Here, we combine life history data with >13,000 behavioural scans and >300 fecal samples collected longitudinally across multiple years from 33 white-faced capuchin monkey females to examine the relationships among behaviour, gut microbiota composition, and reproductive state. We used 16S-based amplicon sequencing and the DADA2 pipeline to analyze microbial diversity and putative functions. Reproductive state explained some variation in activity, but overall resting and foraging behaviours were relatively stable across the reproductive cycle. We found evidence for increases in biotin synthesis pathways among microbes in lactating females, and that relatoe abundance of major phyla among the states was small but significant. Otherwise, gut microbiota structure and estimated functions were not substantially different among reproductive states. These data contribute to a broader understanding of plasticity in response to physiological shifts associated with mammalian reproduction.

## INTRODUCTION

The demands of pregnancy and lactation have been an influential force throughout mammalian evolution. Female mammals experience discrete stages of the reproductive cycle, including cycling, pregnancy, and lactation, but variation across mammalian taxa exists in response to cycling parameters, litter size, birth weight, gestation length, weaning age, weaning mass, and interbirth interval (Gittleman & Thompson, 1988). Lactation is typically the most energetically demanding stage of the reproductive cycle because milk production and other aspects of infant care, incuding infant carrying, require considerable energy above basal metabolic function (Clutton-Brock et al., 1989; Dewey, 1997; Gittleman & Thompson, 1988). Pregnancy is the second most energetically-demanding state, and non-pregnant, non-lactation states (i.e. cycling and non-cycling pauses) are the least energetically costly (Dufour & Sauther, 2002; Serio-Silva et al., 1999). Pregnancy and lactation also introduce increased protein and other nutrient requirements to fuel fetal and infant growth (Dewey, 1997; National Research Council, 2003). Energy requirements typically increase as a fetus develops during pregnancy; after parturition, energy demands continue to increase as the mother produces milk (Ellison, 2003; Emery Thompson, 2013; Villar et al., 1992). As the infant grows and needs more milk combined with larger infant size, energy demand on the mother continues to grow. During the final stages of lactation, once the infant becomes semi-independent in the lead-up to weaning, energy requirements related to infant care decrease (Fig. 1).

**Figure 1.**
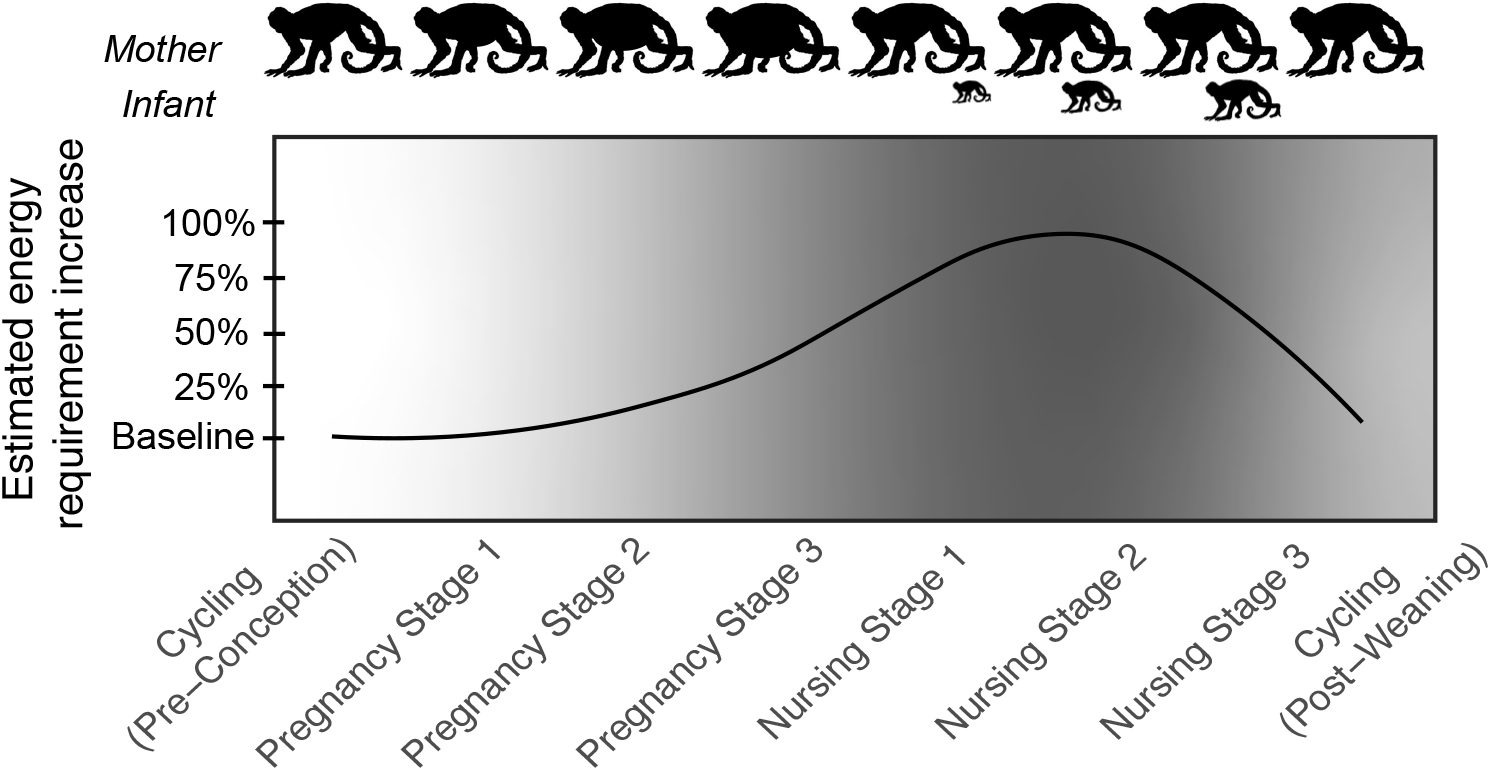
Visualization of estimated increases in energy requirements during the reproductive cycle of a non-human primate. Female primates face a 25% increase in daily energy requirements during gestation, and up to a 50-100% increase during lactation (Key & Ross, 1999).

Mammals vary widely in the length and energy costs of reproduction and have evolved multiple strategies in response. Adaptations include behavioural responses to seasonal fluctuations in food availability. For example, harbour seals *(Phoca vitulina)* and other migratory species travel to specific breeding sites and feeding sites, and exhibit strictly seasonal breeding that is tied to food abundance in their environment (Bowen et al., 2001). For mammals that are not constrained by migratory patterns or strict seasonal breeding, behavioural flexibility—including modulating energy expenditure, foraging rates, and food choice—offers a strategy to mitigate increased energy costs of pregnancy and lactation. Primates, including humans, exemplify these behvioral strategies. While some primates conserve energy during costly reproductive stages by resting for larger proportions of the day (e.g. red-ruffed lemurs [*Varecia rubra*], (Vasey, 2005); green monkeys [*Cercopithecus sabaeus*] (Harrison, 1983)), others increase energy intake, either by foraging for longer periods of the day (e.g. yellow baboons *(Papio cynocephalus*) (Altmann, 2001)) or by increasing their intake rate of foods (e.g. white-faced capuchins [*Cebus capucinus imitator*] (McCabe & Fedigan, 2007)).

Adaptations to the demands of the reproductive cycle also include physiological changes that occur within the mother. For example, changing patterns of fat gain enable females to accumulate fat stores during pregnancy that they can draw from during lactation. Humans typically experience increased fat deposition during pregnancy, even in cases when they are experiencing food stress (Poppitt et al., 1993). Similar results were found in a study of captive bonobos (*Pan paniscus*), in which pregnant females did not lose weight while experiencing caloric restriction (Deschner et al., 2008). Sufficient temporary fat gain during pregnancy supports efficient and healthy development of infants. Too little fat gain may lead to increased periods of lactation and increased interbirth interval (e.g. humans (Lunn et al., 1984)), while too much fat gain during gestation can lead to birth complications (e.g. domestic canines and felines (Fontaine, 2012)).

Research on humans suggests that pregnancy is also associated with changes in gut microbial communities (DiGiulio et al., 2015; Koren et al., 2012; SMID et al., 2018). These changes, which include reduced diversity of microbes, shifts in prominent bacterial phyla associated with energy harvest, and shifts in putative metabolic pathways related to energy absorption are linked to metabolic disease states in non-pregnant individuals. However, in the context of reproductive demands, they may serve an adaptive role in increasing energy harvest from food during times of increased energy need for fetal development and allow for increased fat storage to cope with costs of lactation (Edwards et al., 2017; Koren et al., 2012). In non-human mammals, evidence suggests gut microbiota change during reproduction (e.g. Tibetan macaques *(Macaca thibetana)* (Sun et al., 2021)), and shifts may be hormonally mediated (Mallott et al., 2020). However, other researchers have found that composition and predicted function of gut microbiota remained relatively static throughout pregnancy and into early lactation (Jost et al., 2013). These contrasting findings may be due to differences in study design, methods, sample species and population. Alternatively, they may indicate that the degree to which the gut microbiome can shift during pregnancy is constrained by external factors.

Behavioural and gut microbial changes might interact to address the demands of pregnancy and lactation. However, few studies have combined behavioural and gut microbial data tracked throughout pregnancy and lactation to understand the nuances of how mammals in a wild setting cope with increased energy requirements. Here, we combine behavioural and gut microbial data from a well-studied population of wild non-human primates to examine the strategy or combination of strategies that female primates employ to address the increased energetic costs of pregnancy and lactation. We studied a population of omnivorous, wild white-faced capuchin monkeys that exhibit moderately seasonal breeding. Specifically, we examine adult female monkey responses to changing reproductive stages over the course of 5 years in a seasonal dry forest. We combine a robust data set of >13000 behavioural scans with >300 fecal samples collected from 33 monkeys to study behavioural and gut microbial responses to reproduction in a species that inhabits a dynamic and seasonal ecosystem. Our first aim was to compare activity budgets of white-faced capuchins among and within cycling, pregnancy, and nursing stages. We predict that if capuchins employ an “energy conservation” approach during pregnancy and nursing, then females will rest more in stages of higher energy demand compared to stages of lower energy demand. Conversely, if capuchins employ an “energy maximization” approach during pregnancy and lactation, then females will forage for larger proportions of their day compared to cycling capuchins. Our second aim was to investigate gut microbial changes in female capuchins among reproductive states. We predict that gut microbiota will exhibit characteristics associated with increased capacity for energy harvest during periods of highest energy demand during pregnancy. We also predict that females’ gut microbiota will exhibit an increase in relative abundance of putative metabolic pathways related to energy metabolism and carbohydrate transport during pregnancy. Given the demonstrated potential for ecological and social factors to influence behavioural or gut microbial flexibility in this species, we additionally examine the potential effects of precipitation, temperature, diet, fruit biomass, and dominance rank on activity budget and gut microbial communities.

## MATERIALS AND METHODS

### Field site & study population

We collected samples and behavioural data at Sector Santa Rosa (SSR), located in the Área de Conservación Guanacaste (ACG), in Guanacaste, Costa Rica (10°53’01’’N 85°46’30’’W). Sector Santa Rosa is a mosaic of forest types, including tropical dry forest and small patches of older growth evergreen forest. The ACG experiences two distinct seasons: a hot, dry period from late November to mid-May and a cooler, rainy period for the remainder of the year, during which almost all of the annual rainfall (900 mm-2400 mm) occurs (Melin et al., 2020). Fruit abundance varies throughout the year and estimates of fruit biomass are calculated monthly (Campos et al., 2015; Orkin et al., 2019).

The study population of white-faced capuchin monkeys has been continuously monitored non-invasively since 1983. Female capuchins are philopatric and reach reproductive maturity by 6 years of age. Births are moderately seasonal at Sector Santa Rosa, with 44% of births occurring between May and July each year (Carnegie et al., 2011). Gestation is 157 +/− 8 days and typical inter-birth intervals are 2.5 years (Melin et al., 2020). Lactation lasts for approximately 12 months; in early lactation, infants are almost exclusively dependent on their mothers and are observed nursing frequently (Fragaszy et al., 2004). It should be noted that in other white-faced capuchin populations, the lactation phase can extend to 23 months (Melin et al., 2020).

We collected fecal samples non-invasively from 33 adult females from 4 social groups during multiple sampling bouts in 2014-2016. We collected behavioural data from 33 adult females from the same 4 social groups during multiple sampling bouts in 2016-2018. All animals in the study population are habituated to researcher presence and individually identifiable through physical markings on the face and body. 2016 was the only year in which we collected behavioural records and fecal samples simultaneously.

Pregnancies during the study period were determined via protrusion of the abdomen (visible approximately 8 weeks after conception), and after infants were born we estimated conception dates using 157 days as gestation length. At 15 time points throughout the 5-year study, females exhibited protruding abdomens consistent with pregnancy, but then were later observed with flat abdomens. We characterised these instances as pregnancy loss, though we do not have hormonal data to confirm these pregnancies, which is a limitation associated with this assumption. We determined nursing on an *ad libitum* basis through observations of young monkeys suckling from adult females. Following Bergstrom (2015), we considered females nursing their own infants <12 months of age to be lactating. Juvenile capuchins are occasionally observed suckling after 12 months of age, but it is difficult to determine whether milk is transferred and is less likely. We grouped all non-pregnant, non-nursing females into the category “cycling” following Bergstrom (Bergstrom, 2015).

Studies of humans and non-human primates suggest that energy requirements change throughout pregnancy and lactation (Emery Thompson, 2013). To examine differences that occur *within* each reproductive state, we subset the reproductive states into stages: Cycling (Pre-conception), Pregnancy Stage 1 (early), Pregnancy Stage 2 (mid), Pregnancy Stage 3 (late), Nursing Stage 1 (early), Nursing Stage 2 (mid), Nursing Stage 3 (late), and Cycling (Post-weaning) (Table 1).

**Table 1.** Pregnancy and nursing were divided into three equal stages. Cycling (Pre-conception) consisted of 60 days prior to a conceptive event, and Cycling (Post-weaning) consisted of 60 days post-weaning.

| Reproductive State | Stage | Length |
| --- | --- | --- |
| Cycling (Pre-conception) | -- | 60-0 days before conception |
| Pregnancy | early | 0-53 days post conception |
|  | mid | 54-104 days post conception |
|  | late | 105-158 days post conception |
| Nursing | early | 0-121 days postpartum |
|  | mid | 122-242 days postpartum |
|  | late | 243-365 days postpartum |
| Cycling (Post-weaning) | -- | 0-60 days post weaning |

The reproductive state of each of the 33 adult female capuchins is presented in Fig. 2. Throughout the 2014-2018 study period, 43 infants were born in the study population. Behavioural data collection periods (2016-2018) included portions of or the full duration of 40 of these pregnancies. Of these 40 infants, 26 infants survived to weaning (365 days), and behavioural data collection included portions of all 40 nursing periods and captured transitions from nursing to non-nursing states.

**Figure 2.**
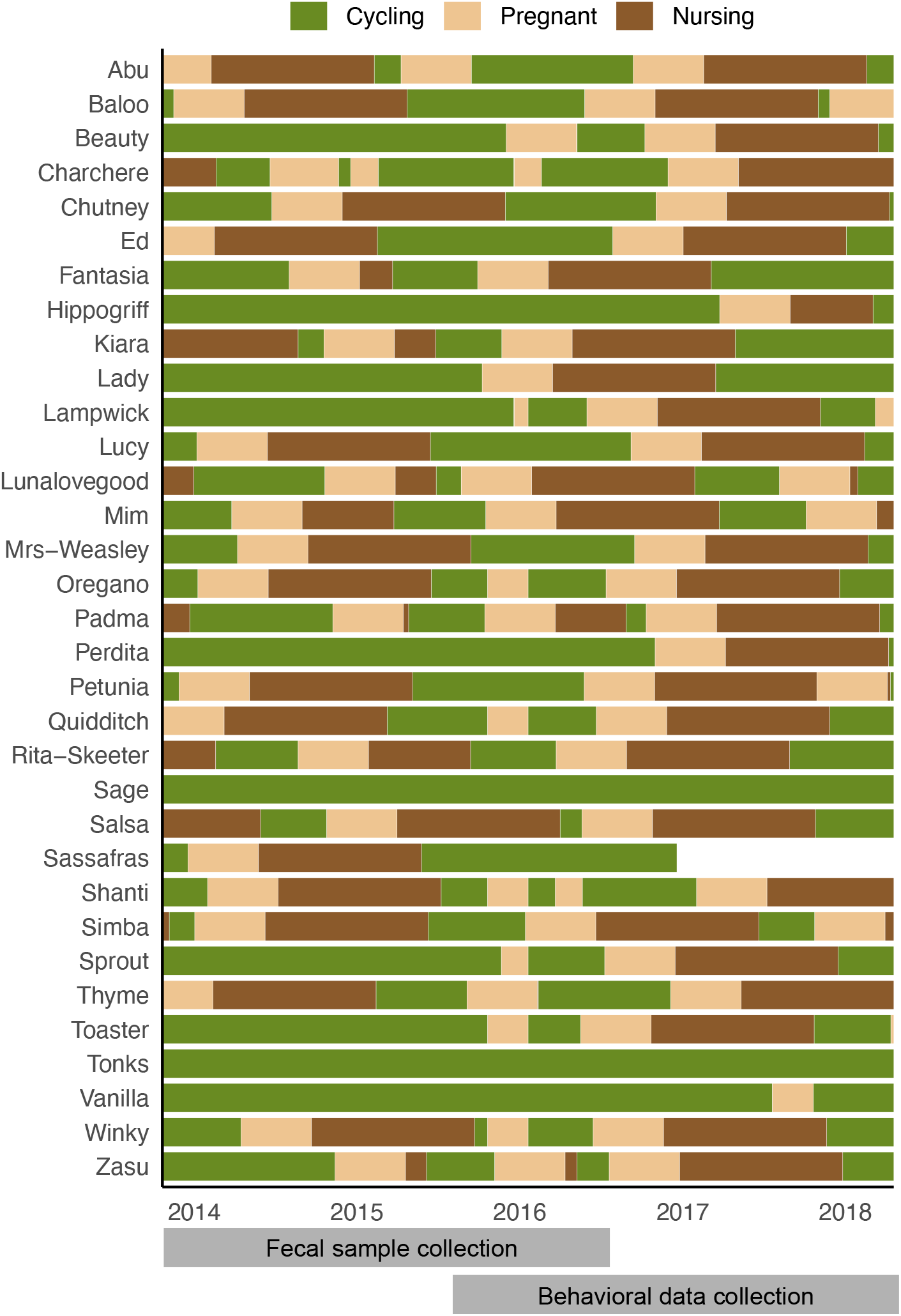
Reproductive states of study individuals observed between April 1, 2014 and June 30, 2018. Fecal samples collection took place between April 29, 2014 and September 27, 2016. Behavioural data collection took place between April 20, 2016 and June 22, 2018. One individual (Sassafras) disappeared from the population in early 2017. Two individuals (Tonks and Vanilla) reached reproductive maturity during the study, but were never observed to be pregnant. Multiple females were observed to be pregnant via protrusion of the abdomen, but were subsequently observed with no protrusion. In these cases the tan (pregnancy) segments are followed by green (cycling) and not by brown (nursing) segments. Nursing segments that are shorter than 12 months represent cases where infants died.

### Daily individual activity budgets

We followed each of the 4 social groups from dawn (05:30) until dusk (18:00) for 4-6 days per month. Individual scans were recorded every 30 minutes on the hour and half hour. During a 10-minute period, we recorded the behavioural state of each individual in the group using an established ethogram (Supplemental Table 1). We chose to use scan sampling instead of focal sampling to determine individual activity budgets because it allows for more evenly distributed data across all individuals, season, and time of day (De Ruiter, 1986; Melin et al., 2018). Inter-observer reliability was tested daily for the first 4 weeks of each sampling period, then weekly or biweekly for the remainder of each period. We collected 13721 individual scans over the course of 222 contact days.

### Behavioural models for activity budget analysis

We fit two generalized linear mixed models (GLMMs) that included reproductive stage as our predictor of interest. For our Resting Model, number of resting scans per day was our response variable, while for our Foraging Model, number of foraging scans per day was our response variable. In each model, we included monkey dominance category, daily maximum temperature (°C), daily rainfall (cm), and mean monthly estimated fruit biomass (kg/ha) as predictor variables as they may influence activity in this population. Ecological variables (e.g. maximum temperature, rainfall, and fruit biomass) were *z*-transformed (i.e. scaled so that each had a mean of 0 and standard deviation of 1) to stabilize the models. We included individual animal identity as a random effect in all models. Sampling effort (i.e. number of scans per animal per rotation) varied due to changing field conditions and stochastic movement and dispersal of group members. We included a log-transformed offset of total scans per animal per day to account for differences in sampling effort. Because our behavioural data are count data and because behavioural scans occur independently, a Poisson distribution with a logit link was designated in all models.

We tested whether our alternative models (fixed and random effects) outperformed the null models (random effects only). Likelihood ratio tests were conducted using the R function ANOVA. To test for multicollinearity between ecological variables a generalized linear model (GLM) was created to determine the variance inflation factor (VIF) (Craney & Surles, 2002). These models were identical to the alternative models above but contained only fixed effects. The resulting VIF measures collinearity in fixed effects. Craney and Surles (2002) suggest that appropriate cutoffs for VIF range from 5-10. All ecological variables had VIF indices below 2.0 and were kept in all models (Supplemental Table 2).

We computed incidence rate ratios using the outputs of our GLMMs to examine the effects of each predictor variable. For categorical variables, the incidence rate ratios represent the ratio of the number of scans recorded in one level compared to the number of scans recorded in another level. For variables with multiple levels (e.g. reproductive stage, dominance), a reference level is selected and other levels are compared to the reference level to contextualise the effects of each level on the response variable—in our case, resting scans or foraging scans. We plotted the predicted outcomes for each reproductive stage using the plot_model function in the R package sjPlot (Lüdecke, 2021).

### Fecal sampling for gut microbiota analysis

We collected fresh fecal samples from study individuals 1-2 times per month within each sampling period in 2014-2016. Once an animal defecated, we immediately collected the feces into a sterile 2mL cryovial using personal protective equipment to minimize contamination. Fecal samples were visually inspected for dietary components, many of which are identifiable by seed shape or arthropod exoskeleton remnants in the feces. Dietary components were recorded to the most specific taxonomic classification possible. Samples were stored on ice in insulated field packs for a maximum of 5 hours before being transferred to a liquid nitrogen shipper (−90 C) for the remainder of the field season. At the conclusion of each sampling season, samples were shipped to the University of Calgary for processing. Samples were collected with permission from the government of Costa Rica from CONAGEBIO (Approval No. R-025-2014-OT-CONEGABIO) and exported under the Área de Conservación Guanacaste permit (DSVS-029-2014-ACG-PI-060-2014). Samples were imported into Canada with permission from the Canadian Food Inspection Agency (Import Permit: A-2016-03992-4). All data collection complied with Costa Rican law and were approved by University of Calgary Animal Care Committee (#AC15-0161).

### Laboratory processing

We used a customized phenol:choloroform-based extraction protocol that included a bead-beading step (Orkin et al., 2019). We purified extracted DNA using an Invitrogen PureLink PCR Purification kit (ThermoFisher Scientific Part No. K310001), after which we combined extractions A and B prior to library preparation. Illumina amplicon sequencing libraries were prepared in for the V4 region of the 16S rRNA gene at the University of Minnesota following Gohl et al. (2016). Libraries were sequenced twice at the University of Calgary to increase reads per sample on an Illumina MiSeq using v2 chemistry.

### Amplicon data preparation

Raw reads were demultiplexed and barcodes and indices were removed using cutadapt (Martin, 2011). We removed ambiguous base calls using the filterAndTrim function in the R package DADA2, removed locus-specific primers using cutadapt, then determined quality profiles using the plotQualityProfile (Callahan et al., 2016). Poor quality bases were truncated again using the filterAndTrim function. Error rates were learned and dereplication was done using learnErrors and derepFASTQ functions respectively. We merged forward and reverse reads to generate amplicon sequence variants (ASVs). Chimeras were removed using the removeBimeraDenovo function in DADA2, and we assigned taxonomies to ASVs using the silva_nr_v132_train_set.fa file. We extracted and sequenced a series of negative lab controls, which were then used to remove probable contaminants in the program decontam (Davis et al., 2018). We then removed uncharacterized phyla, chloroplasts, and mitochondrial sequences.

### Gut microbiota community structure

To explore shifts in gut microbial community structure throughout the reproductive cycle, we computed Chao species richness and Shannon alpha diversity for each sample. We removed 4 samples with Chao1 richness values >400 that were distinctly different that the remaining 304 samples, with Chao1 values ranging from 12-385. Because we sampled individuals multiple times, and because sampling effort across individuals was uneven, we fit linear mixed effects models to examine the relationship between reproductive state and richness and diversity metrics. We included individual identity as a random effect in both models and included rainfall and maximum temperature as ecological predictors and used an alpha of 0.05.

We then removed extremely low-prevalence phyla for the remainder of analysis and filtered out taxa that were not present in at least 5% of samples. Due to sample size constraints, we were not able to divide fecal samples into subsets within reproductive states and therefore proceeded with the categories cycling, pregnant, and nursing. To explore the relationship between reproductive state and gut microbial community dissimilarity within our sample set, we transformed sample counts to relative abundances and then computed Bray-Curtis dissimilarity values using the ordinate function in phyloseq. We visualized beta diversity using non-metric multidimentional scaling (NMDS). We used the function adonis in the R package vegan to run a PERMANOVA to examine predictors of Bray-Curtis dissimilarities in our dataset (Dixon, 2003). In this PERMANOVA, we included reproductive status as our predictor of interest, as well as individual identity, rainfall, and dietary category based on fecal contents, as we suspected these could be related to microbial community dissimilarity.

### Differential abundance

To examine which bacterial taxa were differentially abundant among reproductive states, we agglomerated samples at the genus level, then used the R package DESeq2 to compute variance stabilized counts (Love et al., 2014). We then used Wald tests to determine the log2 fold differences among the reproductive states and used adjusted *P* values (alpha = 0.01) to account for multiple tests. We conducted pairwise comparisons between reproductive states to examine if these transitions are related to gut microbial community structure. We repeated this analysis for bacterial phyla in our dataset to examine courser scale shifts in fecal microbial community structure.

### Estimated metabolic pathways

We used the package PICRUSt2 to estimate metabolic pathways present in our samples using KEGG orthologs (Douglas et al., 2020). We tested for significant dissimilarity in estimated metabolic pathways among the reproductive states using a PERMANOVA including individual identity as a control. We then used the linear discriminant analysis (LDA) effect size method using the LEfSe package (Segata et al., 2011), which identifies the functional metabolic pathways likely to explain differences between reproductive states. We used a logarithmic LDA score of 2 as a cut off for discrimant features, and individual identity was included as a predictor to account for individual variation. All code used for analysis in this study is available at https://github.com/webbshasta/CapuchinReproductionBehaviourMicrobiome.

## RESULTS

### Aim 1 Compare activity budgets of white-faced capuchins among and within cycling, pregnancy, and nursing stages

To visualize overall activity budget shifts across the reproductive cycle, we combined related behaviours (see Ethogram, Supplemental Table 1) into six general categories: Foraging, Resting, Social Affiliation, Social Aggression, Travel, and Other. We calculated proportions of each category per total scans per day (Fig. 3).

**Figure 3.**
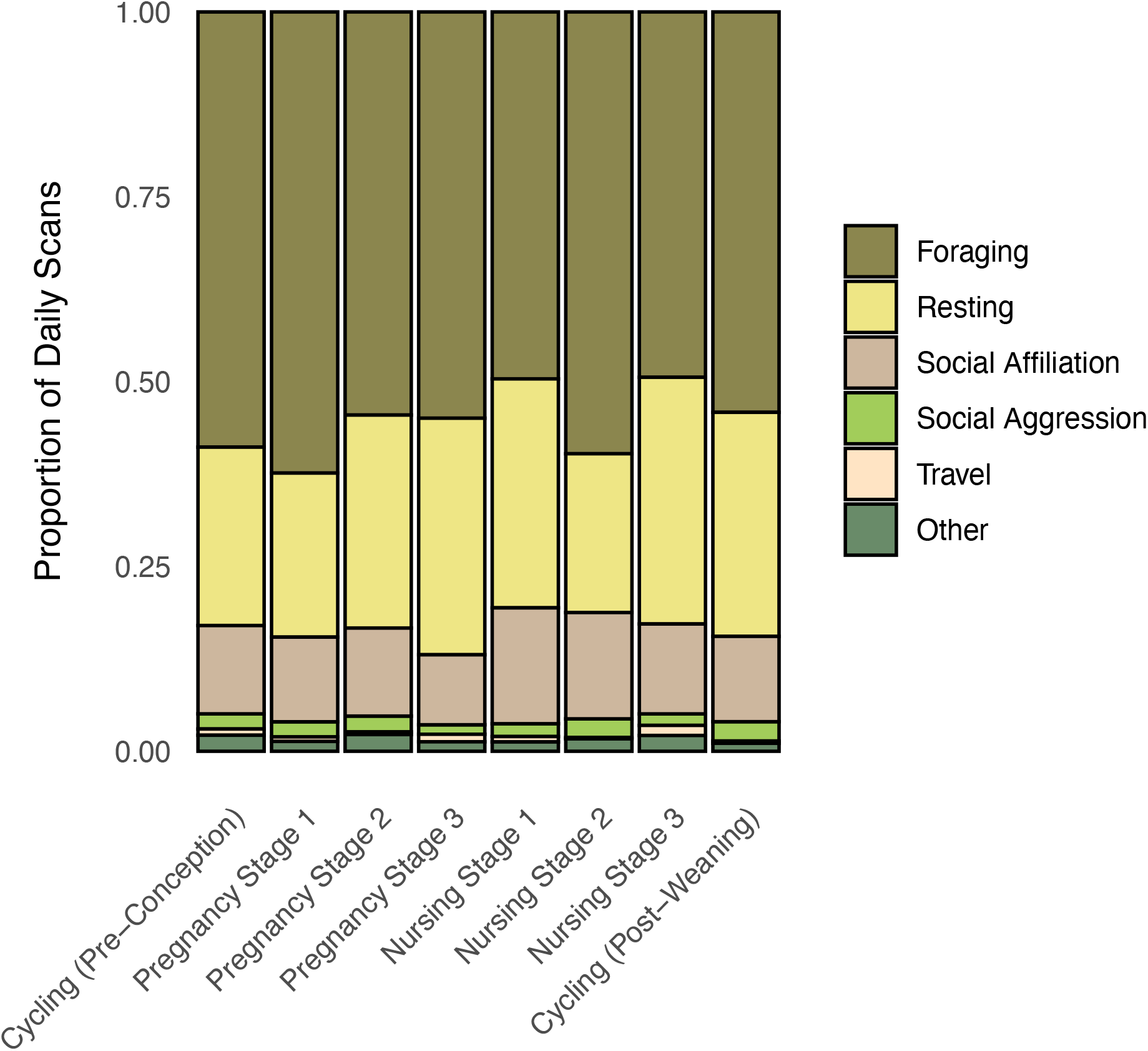
Proportions of daily scans spent in each behavioural category across the reproductive cycle. Daily scans from all individuals were summed for each reproductive state, then divided by total scans per day in that state to determine relative frequency of each behavioural category.

#### Resting activity within and among reproductive states

A generalized linear mixed model of resting activity that included reproductive state outperformed a null model excluding this variable, suggesting some variation in resting behaviour was explained by reproductive stage. High social rank was significantly negatively related to total resting scans (Estimate = −0.13, SE = 0.06, Z-Value = −2.16, p = 0.03), indicating that higher ranking individuals rested less often than lower or mid-ranking individuals. Maximum temperature was significantly positively related to total resting scans indicating that monkeys rested more often in hot temperatures (Estimate = 0.22, SE = 0.02, Z-Value = 10.35, p < 0.001). Incident rate ratios for all predictors are presented in Fig. 4a and values reported in Supplemental Table 2. Predicted counts of resting scans per day are visualized in Fig. 4b and demonstrate that resting increased throughout pregnancy and early nursing, dipped in mid-nursing, and increased again in late nursing. However, variation was minor and we did not find significant pairwise differences among the eight individual reproductive stages.

**Figure 4.**
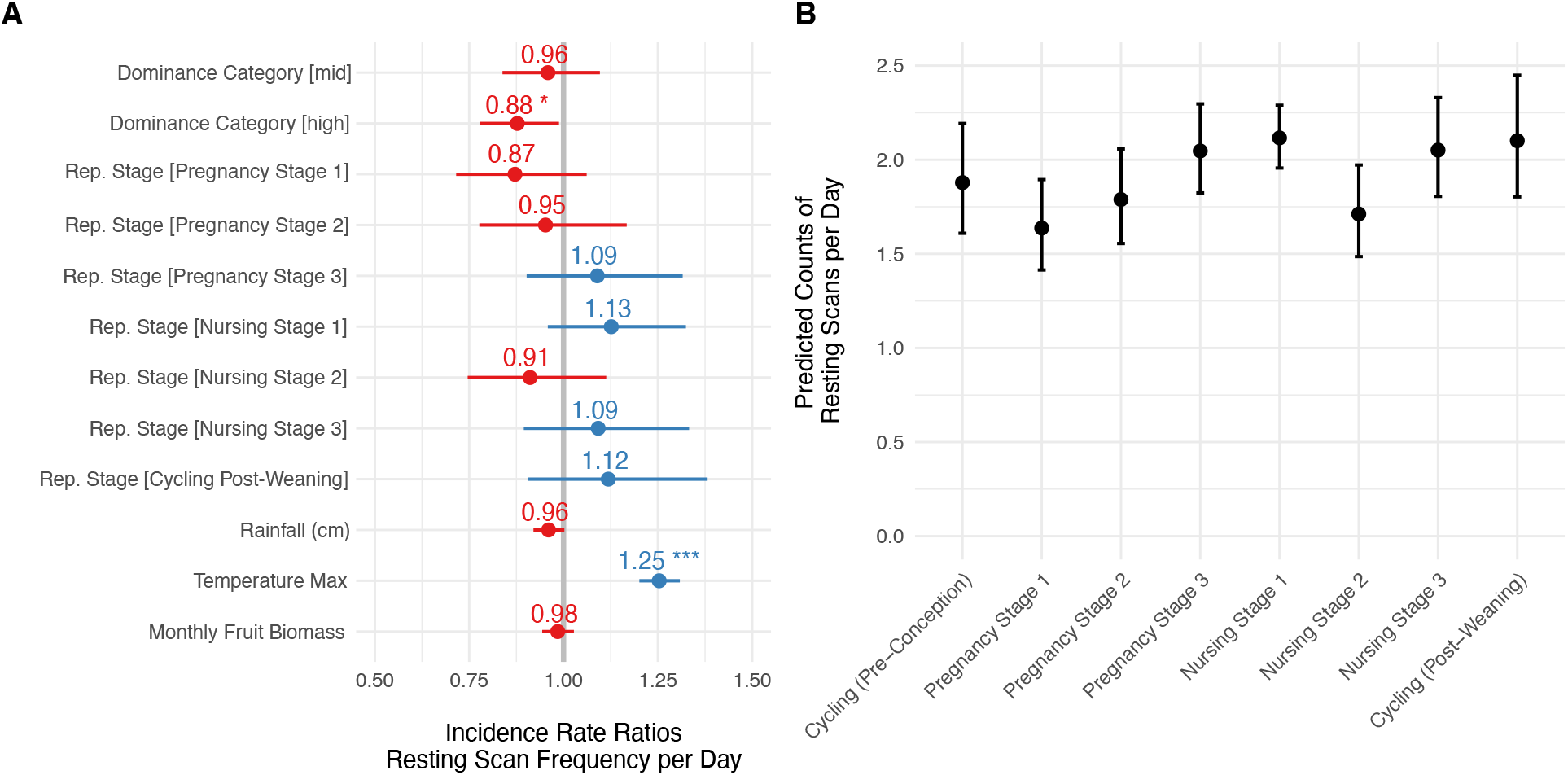
Incidence rate ratios and standard error (**A**) for predictors from a GLMM of resting scans per day. The reference dominance category is low social rank; the reference reproductive stage is cycling (pre-conception). The grey vertical line represents “no effect”. Values to the right of the grey line represent positive effects and values to the left represent negative effects. Significant predictors (p < 0.05) are denoted with asterisks. We plotted the predicted number of resting scans per day for each level of reproductive stage variable (**B**).

#### Foraging activity within and among reproductive states

The GLMM of foraging activity that included reproductive state outperformed the null model. Females in Nursing Stage 1 exhibited significantly fewer foraging scans per day compared to other stages (Estimate = −0.13, SE = 0.05, Z-Value = −2.45, p = 0.01). Ecological variables including rainfall, daily maximum temperature, and estimates fruit biomass were also significantly correlated with foraging scans per day and values are reported in Supplemental Table 2. Incidence rate ratios for all predictors in the model are presented in Fig. 5a and we visualized predicted counts of foraging scans per day in Fig. 5b. These predicted counts suggest that foraging scans steadily decreased throughout pregnancy and into early nursing before increasing throughout late nursing and into post-weaning cycling.

**Figure 5.**
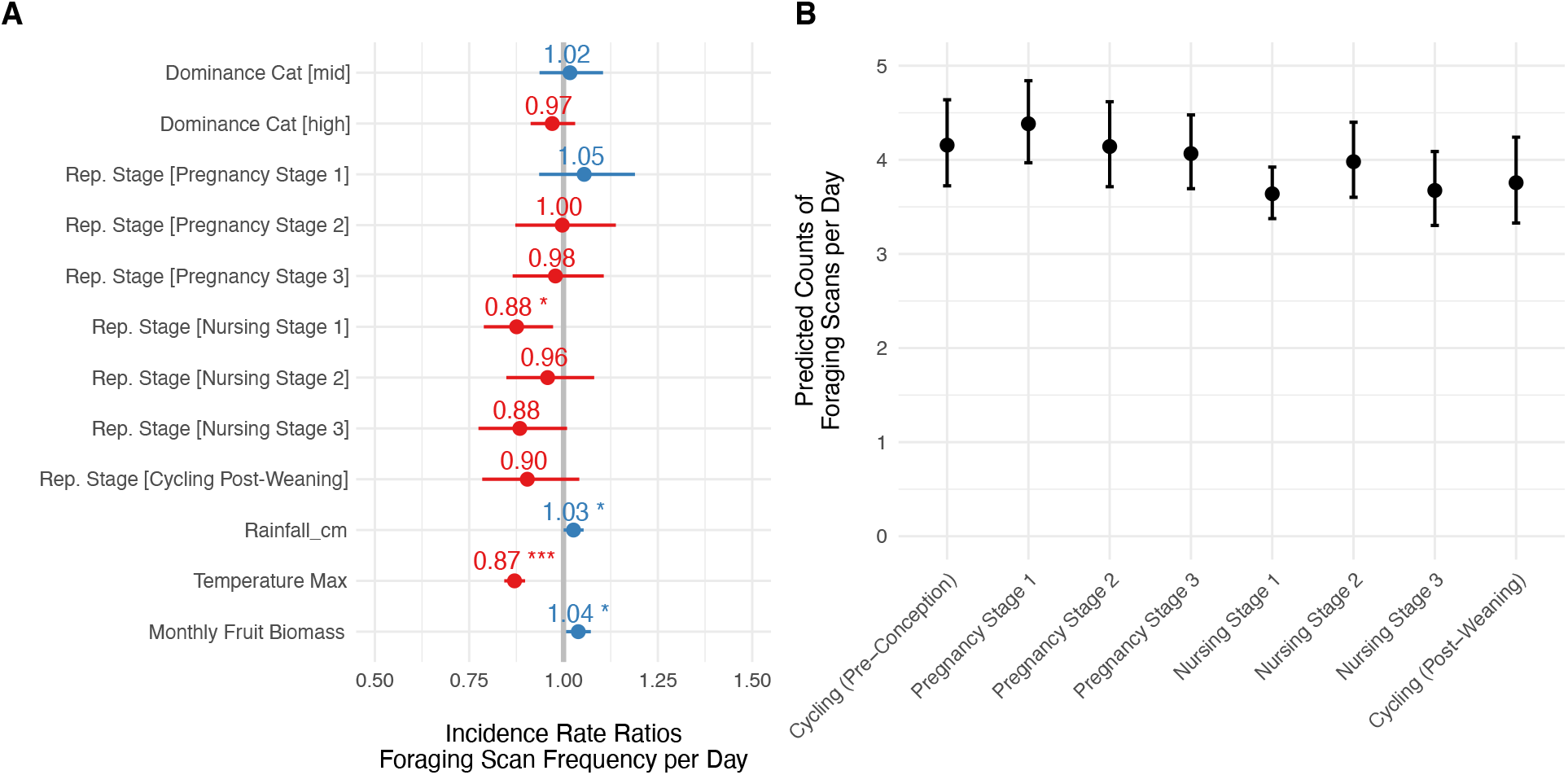
Incidence rate ratios and standard error (**A**) for predictors from GLMM of foraging scans per day. The reference dominance category is low social rank; the reference reproductive stage is cycling (pre-conception). The grey vertical line represents “no effect”. Values to the right of the grey line represent positive effects and values to the left represent negative effects. Significant predictors (p < 0.05) are denoted with asterisks. We plotted the predicted number of resting scans per day for each level of reproductive stage variable (**B**).

### Aim II Investigate gut microbial changes in female capuchins among cycling, pregnant, and nursing states

#### Gut microbial community structure among reproductive states

Reproductive state was not a significant predictor of the Chao1 species richness or Shannon alpha diversity (Supplemental Table 3). Rainfall was significantly negatively correlated with Chao1 richness (Incidence Rate Ratio = 0.89, CI = 0.83 – 0.95, p = 0.001), but no other predictors were significant in either model. Reproductive status was not a significant predictor of gut microbial community dissimilarity (DF= 2, F = 1.275, R^2^ = 0.008, p = 0.163) and samples from the same reproductive state did not cluster distinctly (Fig. 6a). Individual identity accounted for a statistically significant degree of dissimililarity among samples (DF = 28, F = 1.278, R^2^ = 0.11, p = 0.003), as did diet type and rainfall (Supplemental Table 4).

**Figure 6.**
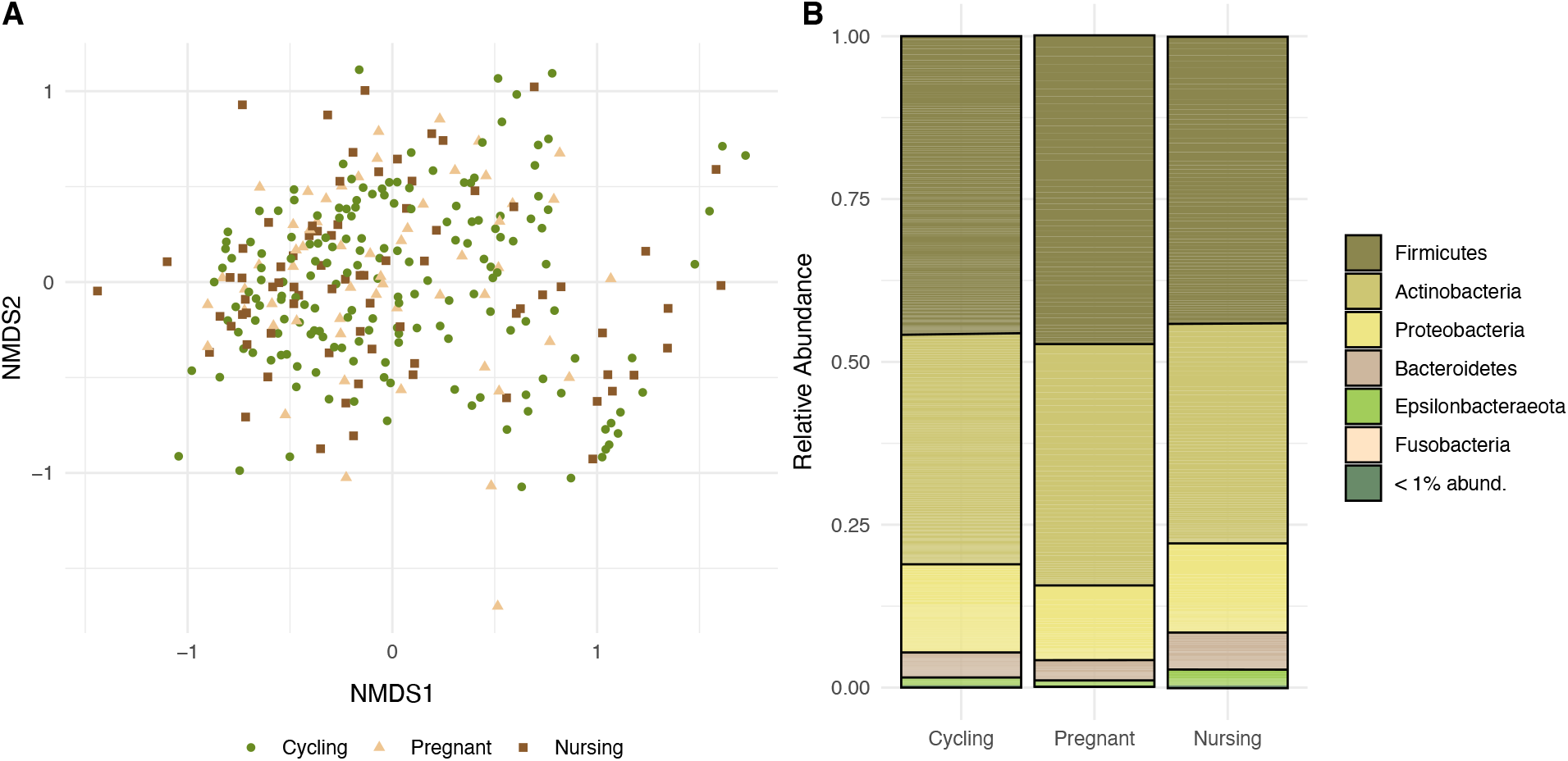
For each sample, Bray-Curtis dissimilarity values were computed and ordinated using non-metric multidimensional scaling (NMDS) (**A**). Reproductive status was not a significant predictor of dissimilarity. Relative abundance of phyla were visualized across reproductive statuses (**B**). Phyla with relative abundances below 0.01 were grouped in the category “<1% abund.”

To investigate other structural changes in the fecal microbial communities among reproductive states, we visualized the relative abundance of phyla across the reproductive cycle, grouping all phyla with relative abundances lower than 1% (Fig. 6b). Pregnant females exhibited a small but significant increase in Firmicutes taxa compared to cycling females (log2 fold change = 0.493, SE = 0.146, p = 0.008). Cycling females exhibited a small but significant increase in Epsilonbacteraeota (log2 fold change = 1.149, SE = 0.293, p < 0.001). We also examined whether specific genera were differentially abundant among the reproductive states and found that cycling females exhibited a significant increase in taxa from the genus *Helicobacter* compared to nursing females (log2 fold change = 1.226, SE = 0.309, p = 0.01).

#### Estimated metabolic pathways remain largely stable among reproductive states

Reproductive status was a significant predictor of estimated metabolic pathway dissimilarity, but the effect was small (PERMANOVA; df =2, F Value = 2.5075, R^2^ = 0.017, p < 0.001). Nursing females were characterized by a significant increase in pathways related to biotin metabolism (Linear Discriminant Analysis; LDA Score = 3.176, p = 0.006), but otherwise metabolic pathways did not differ substantially between reproductive statuses.

## DISCUSSION

We analyzed >13,000 individual scans to explore behavioural responses to reproduction and 308 fecal samples to understand gut microbial community dynamics in a population of reproductively mature female capuchin monkeys. Our main findings are 1) reproductive state explains some variation in activity budget; in particular, foraging decreases significantly in early nursing compared to cycling, though resting and foraging activity remain otherwise stable across the reproductive cycle; 2) reproductive state explains variation in relative abundance of major phyla and putative metabolic pathways; 3) ecological and social variables including maximum temperature, social dominance, and estimated fruit biomass, and individual identity are related to activity and the gut microbiota more strongly than reproductive status.

### Resting and foraging activity within and among reproductive state

During nursing, resting behaviour peaked in early nursing, before decreasing in mid-nursing, and rebounding in late nursing. Foraging behaviour decreased steadily into early nursing, where it was significantly lower in early nursing compared to other stages. We recognize that proportion of scans per day spent in foraging states is an imperfect estimator for amount of food consumed. Nevertheless, the statistically significant drop in foraging behaviour during early nursing may have energy balance implications for females. Capuchins may alter behaviour in other ways to cope with changing energy needs. For example, past research on a small subset of the current study population suggests that lactating females increase feeding rate (McCabe & Fedigan, 2007), though we unfortunately lack the required depth of focal data to test this hypothesis in the current dataset. There might also be underlying metabolic or other physiological changes such as shifts associated with energy sparing that we were not able to capture in the present study that help pregnant and lactating females address energy costs. Even though our sample size of 33 females tracked over multiple years and pregnancies is one of the largest available for wild primates, the pattern we observed in the resting behaviour may be too subtle to reach significance with present sample sizes.

While capuchins are generally considered highly flexible and plastic in response to changes in their environments (Fragaszy et al., 2004), it is likely that both resting and foraging behaviour is relatively constrained in this population and are influenced by social and environmental factors, limiting the potential for flexibility in this domain in response to reproductive state. When food and water resources change drastically from season to season, capuchins alter foraging and ranging behaviours to overlap with available food and water (Campos & Fedigan, 2009). We also see changes in thermoregulatory behaviours; capuchins rest more during the hottest parts of the day in the hotter season of the year, and exhibit seasonal behaviours linked to thermoregulation and water consumption. It is also possible that female capuchins are constrained in altering activity budget due to the pressures associated with group living. White-faced capuchins form cohesive groups, with the exception of emigrant males dispersing to non-natal groups. Females remain the same social group their entire lives (with the rare exception of group fissioning events), and capuchins forage, rest, and travel in close proximity to one another. Pregnant and lactating females may theoretically benefit from resting for longer periods of the day or foraging for longer periods in a particularly productive food patch; however, if a pregnant or lactating female acts independently of the larger social group, she may be increasing risk of predation or encounters with other social groups. Therefore, the ability of females to significantly alter resting or foraging may be constrained by the behavioural choices of the rest of the social group. Future studies examining these constraints on activity budget shifts represent an exciting future avenue of behavioural research.

### Gut microbial community structure among reproductive states

We found mixed support for the prediction that females modulate the gut microbiome to increase energy absorption from food during pregnancy and nursing. In contrast to previous studies in humans that showed a drastic decrease in alpha diversity during pregnancy (Koren et al., 2012), we did not observe significant changes in alpha diversity in pregnant females. Females in cycling, pregnant, and nursing states did not cluster separately in a beta diversity plot, suggesting alternative drivers of community dissimilarity, including individual variation. Overall, female capuchins did not exhibit large gut microbial structural shifts, but we did find that pregnant females exhibited small but significant increases in Firmicutes taxa. At a broad scale, taxa within Firmicutes break down carbohydrates that endogenous host enzymes are unable to metabolize. The ratio of Firmicutes to Bacteroidetes has previously been suggested as a biomarker for increased metabolic activity in the gut (Turnbaugh et al., 2006); however, contrasting reports of this ratio suggest it might not serve as a universal biomarker for increased energy harvest (Magne et al., 2020). Further, the lack of substantial change in relative abundance among bacterial genera in our samples suggest that, in this population, reproductive state is not a critical driver of gut microbial community composition, at least at a broad scale.

Though gut microbiota remained largely stable, nursing capuchins in our population exhibited a significant increase in a biotin metabolism pathway in the gut. Endogenous enzymes as well as microbes can metabolize biotin, which is involved in a broad range of metabolic processes related to fat-, carbohydrate-, and amino acid-utilization in mammals. Biotin deficiency during the reproductive has been linked to tetratogenic effects in pregnancy in mice and humans (Báez-Saldaña et al., 2009). Studies of humans have demonstrated that lactation and pregnancy alter biomarkers of biotin metabolism, and that humans are typically deficient in biotin during pregnancy, though precise requirements of biotin remain unknown (Mock et al., 2002; Perry et al., 2014). Our results suggest that the gut microbiome may play an important role in helping nursing females increase biotin supplementation during fetal growth and infant development.

Previous studies tend to suggest that the gut microbiome shifts considerably throughout the reproductive cycle (Amato et al., 2014; Mallott et al., 2020; Mallott & Amato, 2018). In a recent examination of white-faced capuchin reproductive microbial ecology, Mallott and Amato (2018) examined how gut microbial communities changed across reproductive states in females (n_females_ = 5, n_samples_ = 39) sampled across one year in an aseasonally breeding population of white-faced capuchins. Results suggested that the gut microbiome shifts significantly during the reproductive cycle, including differences in relative abundance of Firmicutes (lower in pregnant versus cycling females) and Actinobacteria (higher in pregnant versus lactating females). Further, the authors found that reproductive state was significantly associated with energy and glycan metabolism (Mallott & Amato, 2018). However, this capuchin population lives in a wet aseasonal forest, with little variation in food and water availability throughout the annual cycle (Mallott et al., 2018). The biome where our present study took place is, by contrast, highly seasonal, with distinct shifts in temperature, water availability and fruit and arthropod abundance (Campos et al., 2015; Mosdossy et al., 2015). In the hot, dry season at Sector Santa Rosa, animals contend with drought and high temperature, while in the rainier, cooler season, these forces are less present. We have observed effects of seasonality on ranging behaviour, activity budget, food choice, and the gut microbiota in this population (Campos et al., 2014; Campos & Fedigan, 2009; Melin et al., 2020; Orkin et al., 2019). Extreme seasonality at the present study site and aseasonality at a different site that is home to the same species of capuchins may have critical implications for our understanding of how flexible and plastic this species is across its home range.

Though capuchins are considered among the most flexible species of platyrrhine primates (Fragaszy et al., 2004; Melin et al., 2020), the reality might be a bit more nuanced. Steig Johnson and Kerry Brown (2018) examined niche breadth in Mesoamerican primates using an ecological niche modeling approach, and found that capuchins were constrained by seasonality of precipitation and temperatures. The temperatures and water availability at our study site are near the limit of suitable conditions for this species, which may explain the lack of flexibility that we see in behaviour and gut microbiota. Understanding how flexibility shifts across a species range and identifying what ecological factors permit or constrain a species’ ability to be flexible is critical to understand not only that species’ history, but also how it might fare as ecosystems face anthropogenic and climate-related changes.

Alternatively, we may be missing the importance of individual variation in response to reproductive states. For example, humans residing in the same population display remarkable differences in response to reproductive demands across our global range; for example, women in the Gambia and Sweden experience high within-group variation in weight gain and energy expenditure throughout pregnancy (Poppitt et al., 1993, 1994) and high inter-individual gut microbiota among members of the same population has been found in humans (Healey et al., 2017; Zhu et al., 2015). We found that individual identity accounted for a significant amount of gut microbial community dissimilarity, which raises questions about individual strategies for coping with reproduction. Further, while activity budgets and amplicon sequencing provide important, thought relatively coarse, data about behaviour and gut microbiota respectively, future research on this population of capuchins could incorporate individual focal data and/or shotgun metagenomic sequencing, both of which would provide a more detailed understanding of capuchin reproductive behavioural and microbial ecology.

How animals respond to the demands of reproduction has important consequences for the viability of offspring, and on a longer term scale, the fitness of a population or species. The intricacies of how animals are able to shift their behaviour and how their gut microbial communities respond to pregnancy and lactation represent a complex but critical area of research. For populations living near the ecological limits of their species ranges, it is especially important to understand the extent to which plasticity in behaviour and gut microbial communities might influence pregnancy outcomes and multi-generational fitness.

## ACKNOWLEDGEMENTS

We thank: Roger Blanco and the Área de Conservación Guanacaste for permitting this research; Saul Cheves Hernandez, Monica Myers, Kelly Kries, Nuria Ferrero, Nile Carrethers, and Ronald Lopez for assistance in the field; and Urs Kalbitzer and Jeremy Hogan for help with analyses. The project that gave rise to these results received the support of a fellowship from “la Caixa” Foundation (ID 100010434) and from the European Union’s Horizon 2020 research and innovation programme under the Marie Skłodowska-Curie grant agreement No 847648 (JDO). The fellowship code is LCF/BQ/PI20/11760004. Research was also supported by The Eppley Foundation for Research (JDO, ADM), the International Center for Advanced Renewable Energy and Sustainability (JDO, ADM), Washington University in St. Louis (JDO, ADM), the Alberta Children’s Hospital Research Institute (ADM, SEW, JDO), the National Sciences and Engineering Research Council (ADM), a Canada Research Chair Tier II (ADM), Sigma Xi (SEW), the American Society of Primatologists (SEW), Alberta Innovates (SEW), and a Vanier Canada Graduate Scholarship (SEW).

## AUTHOR CONTRIBUTIONS

SEW and ADM conceived the study. SEW and REW collected the data. SEW and JDO completed laboratory work. SEW analyzed the data with input from JDO. SEW led the writing of the manuscript. All authors provided feeback on the manuscript at various stages and approved the current submission.

## DATA AVAILABILITY STATEMENT

All data will be made available through Dryad. DOI is forthcoming.

