## Supplemental Tables and Figures for "Activity budget and gut microbiota stability and flexibility across reproductive states in wild capuchin monkeys in a seasonal biome"

**Supplemental Table 1**. Ethogram of behaviors for white-faced capuchin monkeys at Sector Santa Rosa, Costa Rica

| Type of Behavior | Specific Behavior | Code | Description |
|---|---|---|---|
| Foraging | Forage: Insect (Extractive) | EFI | Tearing branches, ripping bark |
| | Forage: Fruit (Extractive) | EFF | Pounding, scrubbing, or breaking open fruits |
| | Forage: Flower | FFL | Feeding on flowers |
| | Forage: Fruit | FFR | Feeding on fruit |
| | Forage: Insect | FIN | Feeding on insects |
| | Forage: Other | FOT | Bromeliad leaves, pith, vertebrates |
| | Forage: Visually | VFO | Actively looking for food, including gleaning insects while moving |
| | Forage: Out of sight | FOS | Monkey is foraging but mouth and/or forelimbs are not visible |
| Resting | Rest (Solitary) | RES | Lying alone, not moving |
| | Rest (Social) | SRE | Not moving, lying down |
| Travel | Travel | TRA | Travel; moving very rapidly, not pausing for foraging or socializing |
| Social Affiliation | Social (Active) | SAC | Monkeys are affiliative; allogrooming |
| Social Aggression | Social (Aggressive) | SAG | Chasing, biting conspecifics |
| Other | Vigilant | VIG | Scanning intently at a long range (not for food) |
| | Drink | DRI | Drink |
| | Excretion | EXC | Excretion of feces, urine, or vomit |
| | Self-Directed | SDI | Auto groom |
| | Play | PLA | Play: biting, chasing, hitting, bouncing, pushing, pulling, etc. |
| | Other | OTH | Inter-group encounter, mobbing predator, sexual behaviour |

**Supplemental Table 2**. Generalized linear models to test variance inflation factor for ecological variables.

| | Generalized Linear Model | | GVIF | Df | GVIF^(1/(2*Df)) |
|---|---|---|---|---|---|
| Variance inflation factor test for ecological variables in Resting Model | glm(TotalRestingScans ~ DominanceCat + RepStateStage + Rainfall_cm + TempMax + MonthlyFruitBiomass + offset(log(TotalScans)), data = dfzGrouped, family = "poisson") | DominanceCat | 1.080238 | 2 | 1.019483 |
| | | RepStateStage | 1.41904 | 7 | 1.025314 |
| | | Rainfall_cm | 1.197425 | 1 | 1.094269 |
| | | TempMax | 1.280846 | 1 | 1.131745 |
| | | MonthlyFruitBiomass | 1.198814 | 1 | 1.094904 |
| | Generalized Linear Model | | GVIF | Df | GVIF^(1/(2*Df)) |
| Variance inflation factor test for ecological variables in Foraging Model | glm(TotalForagingScans ~ DominanceCat + RepStateStage + Rainfall_cm + TempMax + MonthlyFruitBiomass + offset(log(TotalScans)), data = dfzGrouped, family = "poisson") | DominanceCat | 1.080238 | 2 | 1.019483 |
| | | RepStateStage | 1.41904 | 7 | 1.025314 |
| | | Rainfall_cm | 1.197425 | 1 | 1.094269 |
| | | TempMax | 1.280846 | 1 | 1.131745 |
| | | MonthlyFruitBiomass | 1.198814 | 1 | 1.094904 |

**Supplemental Table 3**. Generalized linear mixed models for resting and foraging behaviours. Model results and incidence rate ratios were computed for each prediction.

| Prediction | Generalized Linear Mixed Model | Results | | | | | | | |
|---|---|---|---|---|---|---|---|---|---|
| Females in periods of high energy demand (i.e., pregnancy, nursing) will rest more than females in periods of lower energy demand (i.e., cycling). | TotalRestingScans ~ DominanceCat + RepStateStage + Rainfall_cm + TempMax + MonthlyFruitBiomass + offset(log(TotalScans)) + (1 \| Animal), data = dfzGrouped, family = poisson(link = "log")) | **Predictor** | **Estimate** | **Std. Error** | **Z-Value** | **P-Value** | **Incidence Rate Ratios** | **Confidence Interval** | **P-Value** |
| | | (Intercept) | -1.2988 | 0.08671 | -14.98 | <2e-16 | 0.27 | 0.23 – 0.32 | <0.001 |
| | | DominanceCatmid | -0.04233 | 0.06849 | -0.618 | 0.5365 | 0.96 | 0.84 – 1.10 | 0.537 |
| | | **DominanceCathigh** | **-0.13081** | **0.06053** | **-2.161** | **0.0307** | **0.88** | **0.78 – 0.99** | **0.031** |
| | | RepStateStagePregnancyStage1 | -0.13774 | 0.10057 | -1.37 | 0.1708 | 0.87 | 0.72 – 1.06 | 0.171 |
| | | RepStateStagePregnancyStage2 | -0.04896 | 0.10391 | -0.471 | 0.6375 | 0.95 | 0.78 – 1.17 | 0.638 |
| | | RepStateStagePregnancyStage3 | 0.08571 | 0.09624 | 0.891 | 0.3732 | 1.09 | 0.90 – 1.32 | 0.373 |
| | | RepStateStageNursingStage1 | 0.11913 | 0.08252 | 1.444 | 0.1488 | 1.13 | 0.96 – 1.32 | 0.149 |
| | | RepStateStageNursingStage2 | -0.09301 | 0.10209 | -0.911 | 0.3623 | 0.91 | 0.75 – 1.11 | 0.362 |
| | | RepStateStageNursingStage3 | 0.0879 | 0.10163 | 0.865 | 0.3871 | 1.09 | 0.89 – 1.33 | 0.387 |
| | | RepStateStageCycling_PostWeaning | 0.11204 | 0.10777 | 1.04 | 0.2985 | 1.12 | 0.91 – 1.38 | 0.299 |
| | | Rainfall_cm | -0.04054 | 0.02174 | -1.865 | 0.0622 | 0.96 | 0.92 – 1.00 | 0.062 |
| | | **TempMax** | **0.22587** | **0.02182** | **10.352** | **<2e-16** | **1.25** | **1.20 – 1.31** | **<0.001** |
| | | MonthlyFruitBiomass | -0.01562 | 0.02175 | -0.718 | 0.4726 | 0.98 | 0.94 – 1.03 | 0.473 |
| Females in periods of high energy demand (i.e., pregnancy, nursing) will forage more than females in periods of lower energy demand (i.e., cycling). | TotalForagingScans ~ DominanceCat + RepStateStage + Rainfall_cm + TempMax + MonthlyFruitBiomass + offset(log(TotalScans)) + (1 \| Group/Animal), data = dfzGrouped, family = poisson(link = "log")) | **Predictor** | **Estimate** | **Std. Error** | **Z-Value** | **P-Value** | **Incidence Rate Ratios** | **Confidence Interval** | **P-Value** |
| | | (Intercept) | -0.54966 | 0.060256 | -9.122 | <2e-16 | 0.58 | 0.51 – 0.65 | **<0.001** |
| | | DominanceCatmid | 0.016859 | 0.042153 | 0.4 | 0.6892 | 1.02 | 0.94 – 1.10 | 0.689 |
| | | DominanceCathigh | -0.030515 | 0.031026 | -0.984 | 0.3253 | 0.97 | 0.91 – 1.03 | 0.325 |
| | | RepStateStagePregnancyStage1 | 0.053213 | 0.061225 | 0.869 | 0.3848 | 1.05 | 0.94 – 1.19 | 0.385 |
| | | RepStateStagePregnancyStage2 | -0.003497 | 0.068042 | -0.051 | 0.959 | 1 | 0.87 – 1.14 | 0.959 |
| | | RepStateStagePregnancyStage3 | -0.02176 | 0.062772 | -0.347 | 0.7289 | 0.98 | 0.87 – 1.11 | 0.729 |
| | | **RepStateStageNursingStage1** | **-0.132814** | **0.053359** | **-2.489** | **0.0128** | **0.88** | **0.79 – 0.97** | **0.013** |
| | | RepStateStageNursingStage2 | -0.043159 | 0.061833 | -0.698 | 0.4852 | 0.96 | 0.85 – 1.08 | 0.485 |
| | | RepStateStageNursingStage3 | -0.123121 | 0.067554 | -1.823 | 0.0684 | 0.88 | 0.77 – 1.01 | 0.068 |
| | | RepStateStageCycling_PostWeaning | -0.100933 | 0.072291 | -1.396 | 0.1627 | 0.9 | 0.78 – 1.04 | 0.163 |
| | | **Rainfall_cm** | **0.026021** | **0.013125** | **1.983** | **0.0474** | **1.03** | **1.00 – 1.05** | **0.047** |
| | | **TempMax** | **-0.139173** | **0.016169** | **-8.608** | **<2e-16** | **0.87** | **0.84 – 0.90** | **<0.001** |
| | | **MonthlyFruitBiomass** | **0.038289** | **0.016033** | **2.388** | **0.0169** | **1.04** | **1.01 – 1.07** | **0.017** |

**Supplemental Table 4**. Linear mixed model outputs for richness and alpha diversity among fecal samples across reproductive states and PERMANOVA for Bray-Curtis dissimilarity among fecal samples.

| Study Component | Model Description | Model | Results | | | | | | | |
|---|---|---|---|---|---|---|---|---|---|---|
| Chao1 richness among reproductive states | Generalized linear mixed model with negative binomal distribution | chao1~ ReproductiveStatus + scale(Rainfall) + scale(TemperatureMax) + (1\|INDIVIDUAL), data=metadataFilt) | **Predictor** | **Estimate** | **Std. Error** | **Z-Value** | **P-Value** | **Incidence Rate Ratios** | **Confidence Interval** | **P-Value** |
| | | | (Intercept) | 4.76545 | 0.05274 | 90.356 | <0.0001 | 117.38 | 105.86 – 130.17 | <0.001 |
| | | | Nursing | -0.04053 | 0.09002 | -0.45 | 0.65256 | 0.96 | 0.80 – 1.15 | 0.653 |
| | | | Pregnant | -0.13895 | 0.09505 | -1.462 | 0.14377 | 0.87 | 0.72 – 1.05 | 0.144 |
| | | | **scale(Rainfall)** | **-0.11816** | **0.03632** | **-3.253** | **0.00114** | **0.89** | **0.83 – 0.95** | **0.001** |
| | | | scale(TemperatureMax) | 0.01115 | 0.03721 | 0.3 | 0.76439 | 1.01 | 0.94 – 1.09 | 0.764 |
| Shannon alpha diversity among reproductive states | Linear mixed model with Gaussian distribution | alphadiv~ ReproductiveStatus + scale(TemperatureMax) + scale(Rainfall) + (1\|INDIVIDUAL), data=metadataFilt) | **Predictor** | **Estimate** | **Std. Error** | **T-Value** | **P-Value** | **Incidence Rate Ratios** | **Confidence Interval** | **P-Value** |
| | | | (Intercept) | 2.64774 | 0.0438 | 60.445 | -- | -- | 2.56 – 2.73 | <0.001 |
| | | | Nursing | 0.09473 | 0.07983 | 1.187 | -- | -- | -0.06 – 0.25 | 0.235 |
| | | | Pregnant | 0.08834 | 0.08754 | 1.009 | -- | -- | -0.08 – 0.26 | 0.313 |
| | | | **scale(TemperatureMax)** | **0.07061** | **0.03513** | **2.01** | -- | -- | **0.00 – 0.14** | **0.044** |
| | | | scale(Rainfall) | -0.05154 | 0.03443 | -1.497 | -- | -- | -0.12 – 0.02 | 0.134 |
| Bray-Curtis dissimilarity among reproductive states | PERMANOVA using adonis function R package vegan | distance(psState_filt, method="bray") ~ ReproductiveStatus + INDIVIDUAL + scale(Rainfall) | **Predictor** | **Df** | **Sums of Squares** | **Mean Squares** | **F-Value** | **$R^2$** | **P-Value** | **--** |
| | | | Reproductive Status | 2 | 0.614 | 0.30706 | 1.239 | 0.00789 | 0.186 | |
| | | | **Individual** | **28** | **8.618** | **0.30779** | **1.242** | **0.11071** | **0.005** | -- |
| | | | **scale(Rainfall)** | **1** | **0.711** | **0.71059** | **2.8674** | **0.00913** | **0.003** | -- |
